# Lasso peptides sviceucin and siamycin I have anti-virulence activity and restore vancomycin effectiveness in vancomycin-resistant *Enterococcus* sp. and *Staphylococcus aureus*

**DOI:** 10.1101/2024.07.30.605797

**Authors:** Abdelhakim Boudrioua, Benjamin Baetz, Solenn Desmadril, Christophe Goulard, Anne-Claire Groo, Carine Lombard, Sabrina Gueulle, Marie Marugan, Aurelie Malzert-Freon, Axel Hartke, Yanyan Li, Caroline Giraud

## Abstract

Antibiotic resistance is a major threat to human health and new drugs are urgently needed. Ideally, these drugs should have several cellular targets in pathogens, decreasing the risk of resistance development. We show here that two natural ribosomally-synthesized lasso peptides (LP), sviceucin and siamycin I, (i) abolish bacterial virulence of pathogenic enterococci, (ii) restore vancomycin clinical susceptibility of vancomycin-resistant (VR) enterococci *in vitro* and in a surrogate animal model, and (iii) re-sensitize VR *Staphylococcus aureus*. Mode of action (MoA) analyses showed that they do so by inhibiting the histidine kinases (HKs) FsrC and VanS controlling these phenotypes. Strains resistant to the vancomycin/LP combination were difficult to obtain, and were still fully susceptible to the anti-virulence effect of the LPs, highlighting the advantage of multiple targets. Together with the highly sought-after MoA as HK inhibitors, such properties make these lasso peptides promising candidates for the development of next generation antibiotics.

## Introduction

The alarming increase of infections caused by multidrug-resistant (MDR) bacteria is a main threat for human health worldwide^1^. The World Health Organization (WHO) published a list of the MDR bacteria for which new antibiotics are urgently needed. On this list, vancomycin-resistant enterococci (VRE) as well as methicillin-resistant *Staphylococcus aureus* (MRSA) are classified as “high priority”. These organisms are leading causes of hospital-acquired infections in high-income countries, and a high prevalence of MDR isolates among these strains renders clinical treatments difficult. Vancomycin resistant *Staphylococcus aureus* strains (VRSA) were also reported^2,3^.

*Enterococcus* sp. and *S. aureus* are commensal Gram-positive bacteria^2,4^. *Enterococcus* species are common colonizers of the gastrointestinal tract in humans and other animals^4^. Pathogenic *Enterococcus* species, mainly *E. faecium* and *E. faecalis*, can cross the intestinal barrier and cause bacteremia, endocarditis, and urinary tract infections in immunocompromised patients^5^. *S. aureus* is a commensal of human nasal mucosa. This species is a leading cause of bacterial infections in healthcare and community settings. Skin infections and bloodstream infections occur when mucosal barriers are damaged or owing to invasive medical devices^6^.

Both pathogenic enterococci and *S. aureus* pose a therapeutic challenge due to their ability to form biofilms at the infection sites in addition to their frequent resistance to antibiotics^6,7^. Enterococcal proteases such as the gelatinase GelE and the serine protease SprE are associated with biofilm formation during endocarditis^5^. In *E. faecalis*, biofilm formation is regulated by the Fsr quorum sensing system. The response regulator FsrA and the HK FsrC form a two-component regulatory system (TCS), which is responsible for the quorum sensing-dependent activation of biofilm formation through the activation of *gelE* and *sprE* expression^8,9^.

Vancomycin is the antibiotic of choice to treat enterococci and *S. aureus* infections. Vancomycin inhibits peptidoglycan biosynthesis by binding to N-acetylglucosamine-N-acetylmuramic acid-pentapeptide ending with D-alanyl-D-alanine (D-Ala-D-Ala) dipeptide^10^. Vancomycin resistance occurs following the acquisition of *van* genes, resulting in the synthesis of modified peptidoglycan precursors. The substitution of the last D-Ala of the pentapeptide by D-lactate (D-Lac) greatly lowers the affinity of vancomycin to its target^11^. Several types of *van* clusters have been described. However, VanA- and VanB-types are the most relevant in the clinic^12^. In VanA-type resistance, the D-Lac dehydrogenase VanH converts pyruvate to D-Lac, whereas the ligase VanA forms the dimer D-Ala-D-Lac and the dipeptidase VanX hydrolyses the endogenous D-Ala-D-Ala dimers. The corresponding enzymes in VanB-type resistance are VanH_B_, VanB, and VanX_B_, respectively^11^. The *van* gene cluster is regulated by a TCS consisting of the HK VanS (named VanS_B_ in VanB-type resistance) and the response regulator VanR (named VanR_B_ in VanB-type resistance). In the presence of vancomycin, the kinase VanS undergoes autophosphorylation, which subsequently leads to the phosphorylation of VanR, triggering its dimerization and increasing its affinity for the promoter located upstream of *vanHAX*. In non-inducing conditions, the phosphatase activity of VanS keeps VanR in a dephosphorylated state, preventing the expression of the *van* operon^13^.

The therapeutic efficiency of the few approved antimicrobial agents active against vancomycin-resistant bacteria^14^ seems to be temporary, since resistance to these new drugs has already been described. Therefore, as pointed out by the WHO, new antibiotics or alternative approaches are urgently needed to combat these Gram-positive pathogens with critical clinical importance. The lack of identification of new drugs with novel molecular scaffolds and uncommon mechanisms of action has renewed interest in the investigation of alternative strategies. Ideally, the next generation antimicrobials would interfere with bacterial virulence without inhibiting bacterial growth, which is expected to lower the probability of resistance development because of reduced selective pressure^15,16^. Another promising approach consists in interfering with antibiotic resistance mechanisms to restore the original sensitivity^17^. Such molecules could be used in synergy with conventional antibiotics^18^.

TCSs represent the majority of signaling pathways in bacteria and they control a wide range of behaviors crucial for bacterial adaptation, including virulence, biofilm formation, and antibiotic resistance. Most HKs contain a variable N-terminal sensor domain and a conserved C-terminal cytoplasmic kinase domain^19^. The kinase domain comprises a catalytic ATP binding (CA) and a dimerization/histidine phosphorylation domain. The conservation of the kinase domain allows for the potential design of broad-spectrum antimicrobial agents^20,21^. The toxicity risk of such molecules for humans is likely low, given the fact that HKs are absent in mammals. Pathogenic bacteria often contain a significant number of HKs. For instance, 17 HKs have been identified in the core genome of *S. aureus* strains^22^. This represents multiple possibilities to interfere with bacterial fitness, virulence, and antibiotic resistance, features expected for next generation antibiotics^16^.

We report herein that the lasso peptides sviceucin^23^ and siamycin I^24^ display both anti-virulence and resistance-breaking activity in *Enterococcus* sp. and *S. aureus* through the inhibition of corresponding HKs. They are thus very promising candidates for the development of next generation antimicrobials. Lasso peptides belong to the family of ribosomally synthesized and post-translationally modified peptide, natural products produced by prokaryotes. They display a mechanically interlocked structure composed of an N-terminal macrolactam ring and a C-terminal tail that is held firmly within the ring^25,26^. This compact and rigid scaffold makes them highly stable and excellent protein ligands, leading to enzyme inhibition or receptor antagonism. Sviceucin and siamycin I are among the few representatives of the class I lasso peptides (C1LP), characterized by the presence of two disulfide bonds (Fig. 1a and Fig. S1). They show distinct growth inhibition activity of Gram-positive bacteria: sviceucin is a weak antibacterial whereas siamycin I displays a good activity, notably towards *Enterococcus* sp. and *S. aureus*. In *S. aureus*, siamycin I was shown to bind to lipid II, hence blocking the biosynthesis of peptidoglycan, which is supposed to be the basis of its antibacterial activity^27^. Moreover, sviceucin and siamycin I are known to attenuate the TCS FsrAC-mediated quorum sensing in *Enterococcus*^23,28^, with direct inhibition of the HK FsrC by siamycin I^29,30^. This study extends the activity of these C1LPs to interfere with other TCS-controlled processes, i.e. resistance to vancomycin. Through detailed microbiological, genetic, and biochemical analyses, the molecular target of restoring clinical susceptibility to vancomycin in VRE and VRSA is firmly established to be the HK VanS.

**Figure 1.**
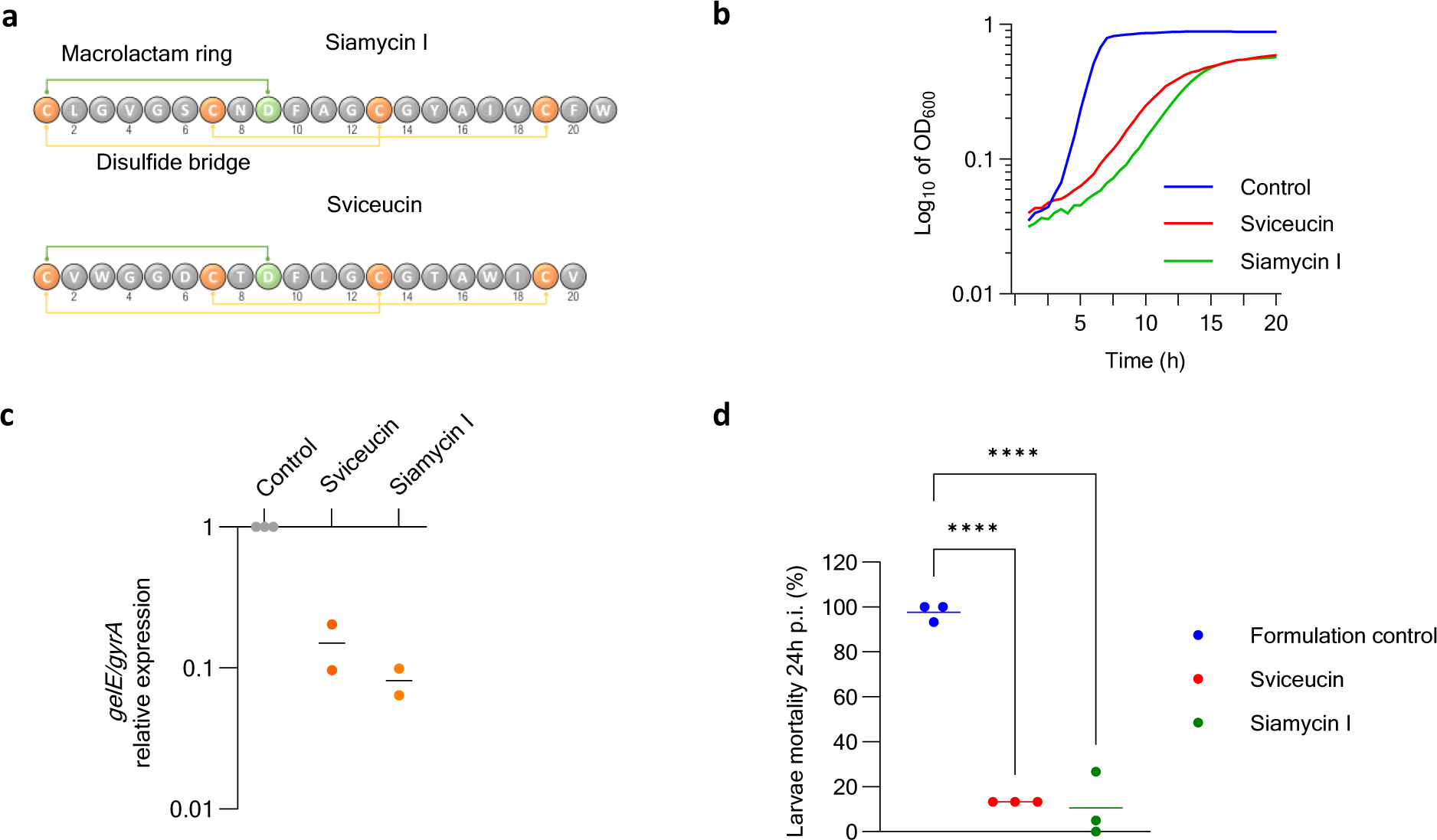
Anti-virulence activity of sviceucin and siamycin I. **a.** Peptide sequences of sviceucin and siamycin I. The macrolactam ring is shown in green. The disulfide bonds are shown in yellow. **b.** Effect of sviceucin and siamycin I on the growth of *E. faecalis* V583. **c.** Effect of sviceucin and siamycin I on the expression of *gelE,* determined by RT-qPCR. **d.** Mortality of *Galleria mellonella* larvae 24 h post-infection (p.i.) with *E. faecalis* V583, pre-treated with either siamycin I, sviceucin or formulation. **** p < 0.0001 (Bonferroni’s multiple comparisons test). Data are averages of three biological replicates.

## Results

### Sviceucin and siamycin I attenuate the virulence of *f. faecalis*

It has been shown that siamycin I binds *in vitro* to the HK FsrC of *E. faecalis* and inhibits its ATPase activity^30^. Therefore, we wondered if siamycin I and the related C1LP sviceucin (Fig. 1a) decreased virulence of *E. faecalis*. Subinhibitory concentrations of siamycin I (2 µM) and sviceucin (10 µM) were used throughout all experiments of this work. At these concentrations, the peptides decreased the growth rate by about 50 % compared to that in the absence of the lasso peptides (Fig. 1b). We first monitored the expression of the Fsr regulated *gelE* gene by RT-qPCR. The peptides were solubilized in a 1:1 water:methanol formulation, and the formulation was used as the control. Both peptides led to a downregulation of expression of *gelE* (Fig. 1c), which is consistent with the previously reported reduction of gelatinase (GelE) activity by sviceucin^23^.

*Galleria mellonella* larvae were then used as a model of *E. faecalis* infection to assess the anti-virulence activity of siamycin I and sviceucin (Fig. 1d). The *E. faecalis* strain V583 was first pre-treated with either C1LP or the formulation and subsequently injected into the larvae. Formulation-treated V583 was highly virulent for the larvae, with an average mortality of 95 % at 24h post-injection. By contrast, bacteria pre-treated with sviceucin or siamycin I were significantly less virulent for the larvae, with mortality rates of 13 % and 10 %, respectively. Similar effects on virulence have been obtained using sviceucin for *E. faecalis* strains MMH594 and OG1RF (Fig. S2).

### Sviceucin and siamycin I restore vancomycin susceptibility

Since sviceucin and siamycin I inhibit signal transduction of the FsrAC TCS *in vivo*, we next wondered if the lasso peptides would also target other TCSs. In *E. faecalis*, the VanRS TCS regulates resistance to vancomycin, an antibiotic of clinical importance. The minimum inhibitory concentrations (MICs) of this antibiotic in combination with the lasso peptides were thus determined for clinically relevant *Enterococcus* and *S. aureus* strains in our in-house collection.

Most interestingly, both lasso peptides re-sensitized VanB-type *E. faecalis* strains to vancomycin whereas only siamycin I was active to restore susceptibility of VanA strains to this glycopeptide (clinical and laboratory standards institute (CLSI) breakpoint > 4 µg/mL). The MIC of vancomycin in strain V583 decreased from 32 to 2 µg/mL in the presence of either one of the two peptides (Table 1), becoming as sensitive as an isogenic strain with a deleted *vanB* gene (V583 Δ*vanB*). This suggests that the effects of the peptides are *van*-operon type dependent. The same activity spectrum of the lasso peptides related to the *van*-operon type was also observed in VRSA strains. Sviceucin re-sensitized only the VanB-type resistant strain, while siamycin I synergized with vancomycin in both VanA- and VanB-type VRSA strains (Table 1). Additionally, we tested the MIC for vancomycin of a vancomycin-intermediate *S. aureus* (VISA) strain in combination with the lasso peptides. VISA strains do not harbor a *van* operon.

**Table 1:** Sviceucin and siamycin I reverse vancomycin resistance. MICs of antibiotic alone, in combination with sviceucin, or with siamycin I.

| Strain | Resistance type | Vancomycin MIC (µg/mL) |  |  |
| --- | --- | --- | --- | --- |
|  |  | Control | Svieceucin | Siamycin I |
| *E. faecalis* V583 | VanB | 32 | 2 | 2 |
| *E. faecalis* JH2-2 08048 | VanB | 64 | 2 | <0.5 |
| *E. faecalis* merz96 | VanB | 32 | 1 | 2 |
| *S. aureus* T-SAR12 VRSA.B | VanB | 64 | 2 | 2 |
| *E. faecalis* 1 231 410 | VanA | 128 | 128 | 2 |
| *E. faecalis* HIP11704 | VanA | 1024 | 512 | <0.5 |
| *E. faecium* 1 231 502 | VanA | 128 | 128 | 2 |
| *E. faecium* 1 230 933 | VanA | 512 | 256 | 2 |
| *S. aureus* T-SAR12 VRSA.A | VanA | >512 | >512 | 2 |
| *E. faecalis* V583 $\Delta vanB$ | *van-* | 2 | 2 | 2 |
| *S. aureus* 16038 (VISA) | *van-* | 4 | 4 | 4 |

The low level of vancomycin resistance (MICs 4 to 8 µg/mL) is due to spontaneous mutations in genes predominantly involved in cell envelope biosynthesis^31^. In the presence of either LP, no reduction of the MIC of vancomycin was observed, further suggesting that the action of these peptides is *van*-operon dependent. Considering the relevance of a vancomycin resistance-breaking activity, we subsequently focused our efforts on (i) providing an in-host proof of the efficacy and (ii) deciphering the underlying MoA of the lasso peptides.

We used *Galleria mellonella* insect larvae to investigate the efficacy of the vancomycin/C1LP combinatorial treatment (Fig. 2). As sviceucin and siamycin I are hydrophobic peptides^23,32^, they were first solubilized in a nanoemulsion previously shown to improve the solubility and the bioavailability of lipophilic drugs^33^. A nanoemulsion with a sufficient drug payload is necessary for the treatment of infected larvae, and this was only obtained for siamycin I, for which it contained 400 µg/mL of the peptide. Larvae were infected with the wild-type strain *E. faecalis* V583, the isogenic vancomycin-sensitive mutant V583 Δ*vanB,* or an equivalent volume of saline solution. At t_90min_ and t_24h_ post-infection (p.i), the larvae were treated with the nanoemulsion alone (referred to as formulation), the nanoemulsion loaded with siamycin I (3.92 µg/larvae; about 18 mg/kg), vancomycin (1 µg/larvae; about 4.6 mg/kg) alone, or a combination of both. Larvae survival was assessed, and statistical analysis was performed at the endpoint t_96h_. The formulation and siamycin I-loaded nanoemulsions did not exhibit toxicity towards the larvae (Fig. 2). 95 % of infected larvae with the WT strain and treated with the formulation were killed 24 h p.i. Siamycin I treatment significantly delayed the killing of the larvae at the beginning of the experiment (p value = 0.0486 at 48 h p.i), although most larvae were also dead by the end of the experiment. This attenuation of the killing rate is likely related to the anti-virulence activity of the peptide. Larvae infected with the WT strain and treated with the combined treatment resulted in a significant improvement of survival in comparison to the condition where larvae were treated with either vancomycin or siamycin I alone. Moreover, the combination therapy resulted in a survival rate similar to that of larvae infected with the V583 Δ*vanB* strain and treated with vancomycin. We therefore provide a proof-of-concept for the potential use of siamycin I as an adjunctive therapy for the treatment of vancomycin-resistant *E. faecalis* in an infected host.

**Figure 2.**
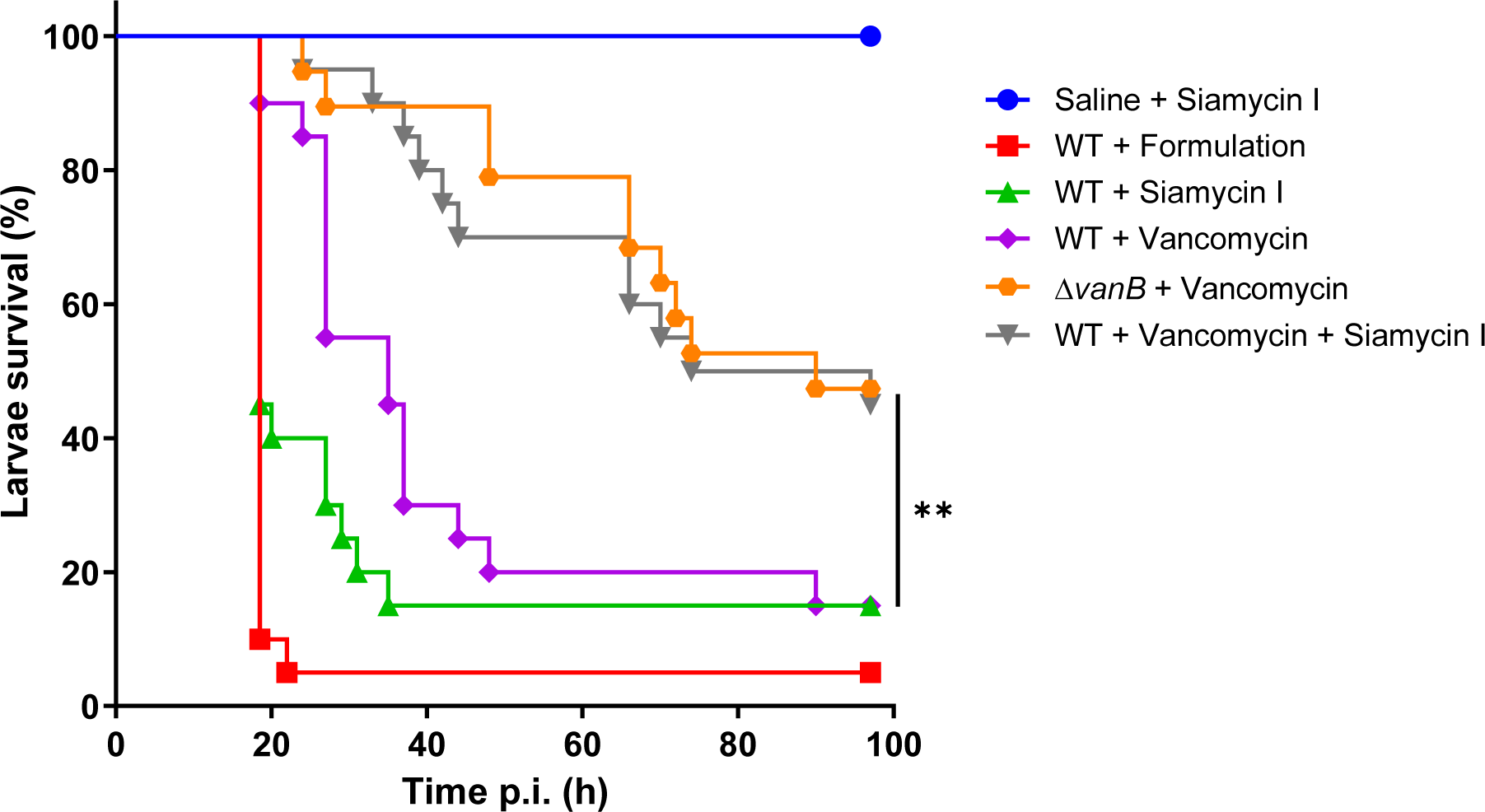
Siamycin I reverse vancomycin resistance. Survival plot of *G. mellonella* larvae infected with 10^6^ CFU of the WT strain, the isogenic Δ*vanB* mutant, or injected with saline solution. Larvae were then treated at t_90min_ and t_24h_ with the formulation, siamycin I, vancomycin or the combination of both. ** p < 0.001 (Logrank test).

### Mutations in VanS_B_ lead to resistance to the vancomycin **/lasso peptide combinatorial treatment**

To gain further insights into the molecular targets of these lasso peptides to reverse vancomycin resistance, resistant clone isolation followed by whole genome sequencing was performed. *E. faecalis* V583, a VanB strain, was subjected to selective pressure in the presence of vancomycin combined with either lasso peptide. Four resistant mutants were isolated when bacteria were plated in the presence of a lethal concentration of combined vancomycin (32 µg/mL) and sviceucin, occurring with an overall frequency of resistance of _∼_10^-8^. By contrast, no resistant mutants could be obtained with lethal concentrations of combined vancomycin (32 µg/mL) and siamycin I. However, by performing serial passages in the presence of 10 µg/mL vancomycin and increasing sublethal concentrations of siamycin I up to a lethal concentration of 5 µM, we were able to isolate five resistant mutants. Three mutants identified from the vancomycin/sviceucin combination have mutations in the locus encoding the F1 ATP synthase, including substitutions in the *atpA* and *atpF* genes and an altered RBS upstream the *atpD* gene. Compared to the wild-type, these mutants were more resistant to vancomycin alone and the vancomycin/sviceucin combination, but still susceptible to the vancomycin/siamycin I combination (Table S1). Four mutants derived from the vancomycin/siamycin I combination have either point mutations or frameshift in the *ccpA* gene encoding the catabolite control protein A. These mutants were resistant to the effects of both C1LPs in combination with vancomycin (Table S1). Interestingly, two mutants, named Svic1 and SiaA and isolated respectively from vancomycin/sviceucin and vancomycin/siamycin I combinations, have a direct link to vancomycin resistance. Genome sequencing revealed that both have point mutations in the ATPase CA domain of VanS_B_, leading to a S402I and a D398E substitution, respectively (Fig. 3a-c). Growth of these mutants was very similar to that of the wild-type strain. Both Svic1 and SiaA exhibited a 4-fold increase of the MIC of vancomycin, reaching 128 µg/mL regardless of the presence of sviceucin (Table 2). In the case of the combination of vancomycin/siamycin I, the SiaA mutant showed partial resistance while the Svic1 mutant remained sensitive. To provide evidence of the causal effect of S402I and D398E substitutions in VanS_B_ on resistance to the lasso peptides, we introduced both mutations individually into a wild-type V583 background. The resistance profiles of V583 *vanS_B_*^S402I^ and V583 *vanS_B_*^D398E^ were similar to the mutants Svic1 and SiaA, respectively, indicating that the resistance phenotypes of Svic1 and SiaA are due to the identified mutations in VanS_B_. Additionally, deletion of *vanB* from the resistant mutant Svic1 led to loss of vancomycin resistance, arguing that the increase of the MIC of vancomycin in Svic1 and SiaA mutants occurs exclusively through *van* resistance genes. Our results strongly support that the histidine kinases VanS and VanS_B_ are direct targets of sviceucin and siamycin I.

**Figure 3.**
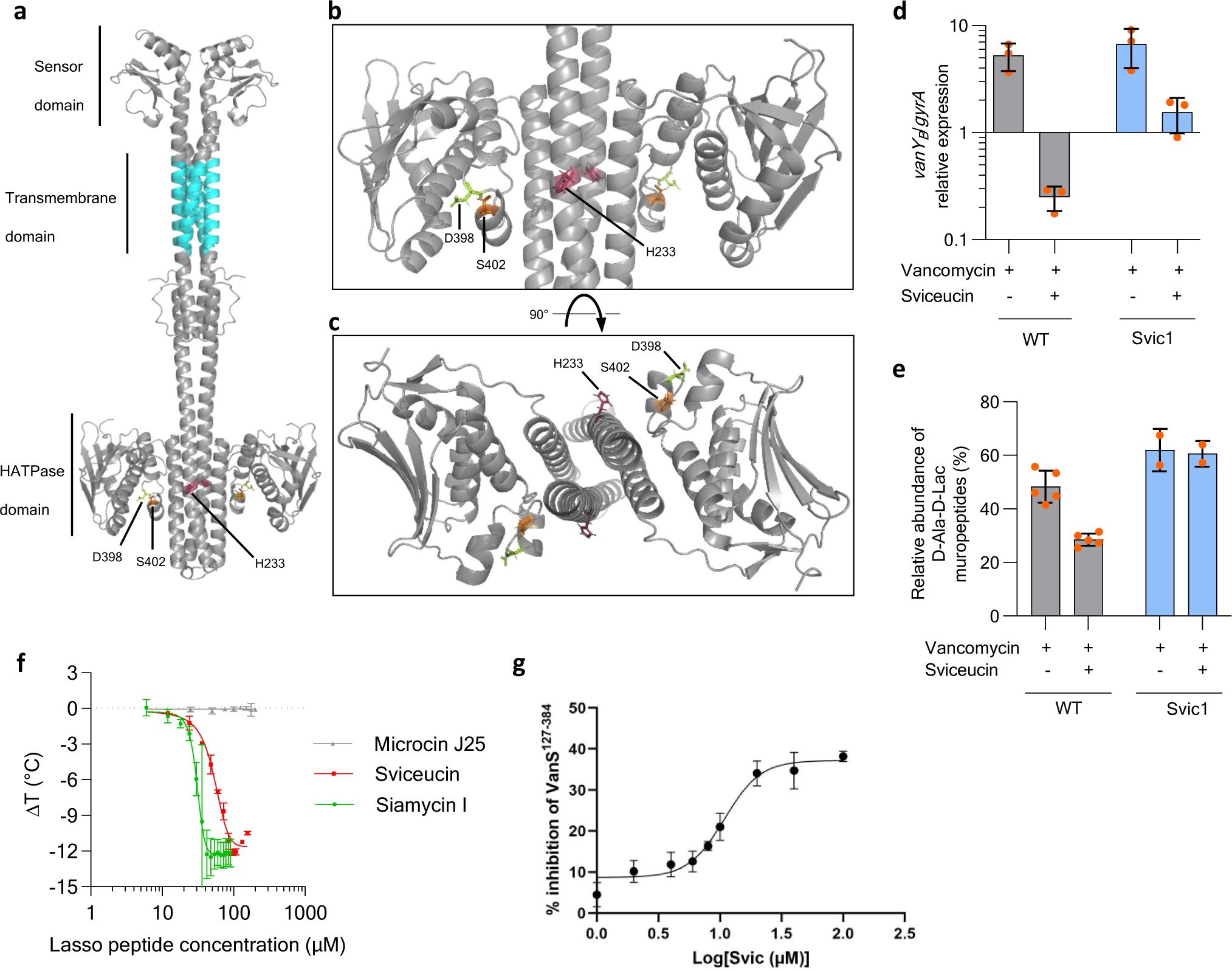
Characterization of VanS/VanS_B_ as the molecular target of sviceucin and siamycin I. **a-c.** Model of the full-length VanS_B_ dimer generated using AlphaFold 1. D398E and S402I mutations and the conserved H233 are mapped on the structure (a). A zoom into the H-ATPase domain (b) and top view of the H-ATPase domain (c), are shown. **d.** Relative expression of *vanY_B_* in presence of vancomycin (1 µg/mL) or vancomycin and sviceucin in *E. faecalis* V583 and its mutant Svic1. **e.** Relative abundance of D-Ala-D-Lac muropeptide in presence of vancomycin (2 µg/mL) or vancomycin and sviceucin in *E. faecalis* V583 and its mutant Svic1. **f.** Differential scanning fluorimetry analysis of VanS-lasso peptides interactions. ΔT of VanS^127-384^ in presence of increasing concentrations of sviceucin, siamycin I or microcin J25. **g.** Dose-response curve of VanS^127-384^ inhibition by sviceucin (IC_50_ 10.8 ± 1.5 µM, nonlinear regression fitting R^2^ = 0.93).

**Table 2.** MICs (µg/mL) of vancomycin of Svic1 and SiaA resistant mutants.

| Strain | Control | Sviceucin | Siamycin I |
| --- | --- | --- | --- |
| V583 | 32 | 2 | 2 |
| *Svic1* | 128 | 128 | 2 |
| *Svic1* $\Delta vanB$ | 2 | 2 | 2 |
| V583 *vanS<sub>B</sub>*<sup>S402I</sup> | 128 | 128 | 2 |
| *SiaA* | 128 | 128 | 32 |
| V583 *vanS<sub>B</sub>*<sup>D398E</sup> | 128 | 128 | 16 |

### Sviceucin leads to a decrease in *van* operon expression

Based on the above results, sviceucin and siamycin I are expected to reduce or block the expression of the *van* operon upon vancomycin induction in the wild-type strain V583 but not in the Svic1 and SiaA mutants. To decipher this part of the mechanism, we chose to focus on sviceucin and the sviceucin-resistant mutant Svic1. The expression of *vanY_B_* was monitored by RT-qPCR when bacteria were cultured with vancomycin (1 µg/mL) alone or in combination with sviceucin. As expected, *vanY_B_* expression was strongly induced by vancomycin alone in both wild-type and Svic1 strains (Fig. 3d). In the presence of sviceucin, *vanY_B_* expression was reduced by a factor of 21 in the wild-type strain upon vancomycin induction, suggesting that sviceucin interferes with the signal transduction of VanR_B_S_B_. By contrast, sviceucin led to only a 4-fold reduction of *vanY_B_* expression in the Svic1 strain. The *vanY_B_* expression level in the Svic1 mutant was 6.2 times higher than in the wild-type strain. This difference may explain the sviceucin-resistant phenotype of the Svic1, mutant in which, conversely to the wild-type, a sufficient expression level of *van* genes is reached to allow synthesis of modified peptidoglycan precursors in order to counter the action of vancomycin.

### Sviceucin leads to a decrease in the percentage of muropeptides ending in D-Ala/D-Lac

To verify the latter hypothesis, we investigated the effect of sviceucin on peptidoglycan biosynthesis in the presence of vancomycin by quantifying the muropeptides ending in D-Ala-D-Ala and D-Ala-D-Lac in the wild-type and Svic1 strains by liquid chromatography coupled to mass spectrometry (LCMS) (Fig. 3e). When induced with vancomycin alone (2 µg/mL), the percentage of muropeptides D-Ala-D-Lac reached 48.3 % and 62.0 % in the wild-type and the Svic1 mutant, respectively, which is consistent with the increased MIC of vancomycin in Svic1. Importantly, the presence of sviceucin resulted in a decrease of the percentage of muropeptides D-Ala-D-Lac (28.5 %) in the wild-type strain, while that of the Svic1 mutant remained unchanged. Taken together, our data confirm that sviceucin breaks *van* operon-mediated vancomycin resistance by affecting the signal transduction of VanRS, leading to a decreased expression of the *van* operon and, subsequently, a decreased incorporation of the modified muropeptides D-Ala-D-Lac into the peptidoglycan. This should also hold true for siamycin I.

### Sviceucin and siamycin I directly interact with the Van histidine kinase

Given that the resistant mutants harbor mutations located in the ATPase domain of VanS_B_, this suggests that the peptides directly interact with this HK. This hypothesis was tested via genetic and biochemical experiments. First, a system allowing VanS-independent constitutive expression of the *van* operon was generated. It has been shown that the conserved aspartate residue D53 at the N-terminal receiver domain of VanR_B_ is the phosphate acceptor following phosphorylation by P-VanS_B_^34,35^, and that substituting the conserved aspartate by a glutamate residue mimics a phosphorylated response regulator (P-VanR_B_)^36^. We introduced a copy of a *vanR_B_* variant gene encoding the phosphomimetic modification D53E into the chromosome of a vancomycin-sensitive V583Δ*vanS_B_* background strain (V583 Δ*vanS_B_ malT*::*vanR^D53E^*). In this strain, the MIC of vancomycin (32 µg/mL) was similar to that of the wild-type strain, indicating that the VanS_B_-independent constitutive expression of the *van* operon was successfully elicited (Table 3). In contrast to the wild-type strain, sviceucin and siamycin I did not suppress the resistance to vancomycin, suggesting that the lasso peptides act upstream of VanR_B_.

**Table 3.** Effect of the phosphomimetic modification VanR^D53E^ on the MICs (µg/mL) of vancomycin.

| Strain | Control | Sviceucin | Siamycin I |
| --- | --- | --- | --- |
| V583 | 32 | 2 | 2 |
| V583 $\Delta vanS_B$ | 2 | 2 | 2 |
| V583 $\Delta vanS_B$ *malT::vanR*<sup>D53E</sup> | 32 | 32 | 32 |

To further support this, we investigated the origins of the different activity spectra of sviceucin and siamycin I between VanA- and VanB-type vancomycin resistance. We first replaced *vanS_B_* by *vanS* in the VanB-type strain *E. faecalis* V583. VanS-VanR_B_ strain was sensitive to vancomycin (MIC = 2 µg/mL), probably because of an impaired signal transduction between VanS and VanR_B_. To bypass this issue, we replaced the *vanR_B_S_B_*-P*_vanYB_* fragment by the *vanRS*-P*_vanH_* fragment from a VanA-type operon, resulting in a VanA-type transduction system fused to VanB-type resistance genes. In this strain, the phenotype of a VanA-type strain was restored, meaning resistance to sviceucin while remaining sensitive to siamycin I (table 4). Furthermore, the replacement of the *vanB* gene encoding the D-Ala:D-Lac ligase by its VanA-type homolog *vanA* did not affect the sensitivity of the strain towards sviceucin, confirming that the activity spectrum of the peptides is linked to the TCS and not the resistance genes. Taken together, these results confirm that sviceucin and siamycin I interfere with the signal transduction of the TCS at the level of the HK VanS_B_/VanS.

**Table 4.** Effect of the signal transduction system on the spectrum of activity of sviceucin and siamycin I on vancomycin MIC (µg/mL).

| Strain | Control | Sviceucin | Siamycin I |
| --- | --- | --- | --- |
| V583 | 32 | 2 | 2 |
| V583 $\Delta vanS_B$-*vanR<sub>B</sub>::vanS-vanR* | 32 | 32 | 2 |
| V583 $\Delta vanB::vanA$ | 32 | 2 | 2 |

To provide evidence for a direct interaction, the cytoplasmic domains of the HKs were recombinantly produced in *E. coli*. Despite numerous efforts, only the C-terminal His_6_-tagged VanA-type VanS^127-384^ could be obtained as a soluble protein. The peptide-protein interactions were analyzed by Differential Scanning Fluorimetry (DSF). The affinities of siamycin I, sviceucin, and an unrelated lasso peptide microcin J25^37^ for VanS^127-384^ were determined by monitoring the changes of the melting temperature (Tm) of the polypeptide (Fig. 3f). While microcin J25 did not affect the Tm of VanS^127-384^, sviceucin and siamycin I caused a dose-dependent reduction of its thermal stability, with an equilibrium dissociation constant (K_d_) of 52±1.6 µM and 38±1.0 µM, respectively. We therefore conclude that sviceucin and siamycin I specifically interact with the VanA-type HK VanS. Next, we investigated whether this interaction leads to inhibition of the autophosphorylation activity of the HK, using a continuous assay based on the detection of ADP^38^. Indeed, sviceucin inhibited VanS^127-384^ with IC_50_ of 10.9±1.5 µM (Fig 3g). By contrast, microcin J25 had no effect on VanS^127-384^ (Fig. S3). Unfortunately, we were unable to determine the IC_50_ for siamycin I since the peptide made the enzymes precipitate under our assay conditions. Given that the CA domains in HKs are highly conserved^21^ and that sviceucin and siamycin I share the same 3D structure (Fig. S1), it is likely that these C1LPs also target the VanB-type kinase VanS_B_.

Collectively, these data confirm that the molecular basis of restoring vancomycin sensitivity by these C1LPs is the direct inhibition of Van HKs. It is worth noting that sviceucin and siamycin I have distinct spectra towards VanA- or VanB-type vancomycin resistance (Table 1), which could, at least in part, be due to the higher affinity of siamycin I for VanS.

## Discussion

Bacteria use TCSs to control a wide variety of physiological responses, including virulence, biofilm formation, competence, and antibiotic resistance, notably in the context of host infection. Interference with these processes likely does not impact bacterial viability, hence reducing the risk of resistance appearance, which is imposed by the selection pressure. Therefore, targeting TCSs represents a promising strategy to combat the antibioresistance crisis. To our knowledge, sviceucin and siamycin I are the only natural products known to target both virulence and vancomycin resistance in major Gram-positive pathogens via inhibition of related HKs. Previous reports and our study show that the C1LPs interfere with FsrC in *E. faecalis*, leading to an attenuation of virulence in a *G. mellonella* infection model. The anti-virulence effect can be extrapolated to *S. aureus* because one of its master virulence regulator, AgrC, is a close homolog of FsrC^39^. This work also establishes that C1LPs inhibit directly VanS, which is the molecular basis of reverting vancomycin resistance. The effectiveness of combined vancomycin/C1LP treatments to restore clinical susceptibility of VRE and VRSA to vancomycin has been proved in the *G. mellonella* infection model. Such an elegant dual mechanism of C1LPs to kill two birds with one stone stands at an immense advantage, as having multiple clinically-relevant targets further decreases the risk of simultaneous developments of resistance to all actions of these molecules. In this regard, siamycin I seems particularly interesting. Siamycin I has another direct target, lipid II, an essential precursor for cell wall biosynthesis in Gram-positive bacteria, and the inhibition of Lipid II led to compromised cell growth^27^.

Breaking antibiotic resistance is a validated therapeutic strategy, as exemplified by the clinical use of β-lactam antibiotic and β-lactamase inhibitor combination^40^. Similar approaches should also be successful to potentiate the use of vancomycin. Being the first-line antibiotic for the treatment of MRSA, β-lactam resistant enterococci, and *Clostridioides difficile* infections^2,3,41,42^, vancomycin encounters resistance frequently in enterococci, especially *E. faecium*, but is hitherto rare for MRSA isolates. In *C. difficile*, vancomycin resistance genes are present and have been associated with elevated vancomycin MICs^43^. High-level vancomycin resistance of *C. difficile* would have dramatic clinical consequences. Therefore, drugs that counteract the vancomycin resistance mechanism would be immediately applicable for VRE infections, but could also be used as precautionary drugs for the treatment of other resistant strains, such as *C. difficile*. Previous studies reported several molecules disrupting the TCS VanSR^14,44^, including a synthetic antimicrobial peptide^45^. However, their molecular mechanisms remained elusive as no direct evidence has been provided that they directly inhibit the TCS VanSR.

To understand the molecular details of how the C1LPs suppress vancomycin resistance, *E. faecalis* mutants resistant to vancomycin in the presence of either sviceucin or siamycin I were isolated. The only overlap between the two combinations were point mutations in VanS. Genetic and biochemical characterization showed C1LPs interact directly with VanS and inhibit its autophosphorylation activity. This leads to blockage of *van* gene expression and subsequent reduction of the incorporation of the modified pentapeptides into the peptidoglycan. Among other spontaneous resistant mutants, mutations have been mapped in the (F_1_F_0_) H+ ATPase (F-ATPase) operon or in the *ccpA* gene, but none of these mutants were shared by the two different combinations. The F-ATPase mutants were specifically resistant to sviceucin, whereas *ccpA* mutants were resistant to both C1LPs. In both cases, introduction of the corresponding mutation in a wild-type background conferred vancomycin resistance in the presence of the C1LPs, although effects on general fitness are expected for such mutants. The F-ATPase functions in lactic acid bacteria, including *E. faecalis*, as a proton pump to maintain intracellular pH homeostasis^46^. In *S. aureus*, ATP synthase mutants show altered growth kinetics and biofilm formation, and are more susceptible to antimicrobial peptides^47,48^. By comparison, CcpA is a global regulator of genes encoding activities for the transport and catabolism of secondary substrates in Gram-positive bacteria^49^. Knock-outs of *ccpA* have pleiotropic effects on the physiology of enterococci and staphylococci, including reduced biofilm formation, antibiotic resistance, and virulence^50,51^. Although not directly linked to the *van* operon, these mutations could provide insights into the interplay between vancomycin resistance and bacterial physiology, which in turn could inspire novel resistance-breaker strategies^40^. Remarkably, these mutants are all still fully susceptible to the anti-virulence activity of the C1LPs, as shown by the *G. mellonella* infection assay (Fig. S4), supporting strongly that developments of resistance to one action of the C1LPs do not induce resistance to other activities of the C1LPs.

Siamycin I and sviceucin have a high structural homology^23^ but differ considerably in several aspects. For both anti-virulence and anti-vancomycin resistance activities, sviceucin requires an effective concentration 5-fold higher than that of siamycin I. Moreover, siamycin I acts on VanA- and VanB-type vancomycin resistant strains, whereas sviceucin is only active on VanB-type strains. Surprisingly, *in vitro* characterization using the soluble cytoplasmic domain of VanS showed that siamycin I only has a slight higher affinity (2-fold) for VanS. The discrepancy of *in vitro* assays and the whole-cell phenotyping could be attributed to: (i) a truncated cytoplasmic domain may not reflect the functioning of a full-length HK, which involves the sensor domain and interactions with the response regulator; (ii) transport into the cytoplasm and subsequence availability are different for these peptides; (iii) binding of siamycin I to lipid II, a membrane-bound precursor for cell wall synthesis, may somehow favor the interaction between this peptide and VanS.

In conclusion, this study identifies the C1LPs as promising HK inhibitors for the development of next generation antibiotics. Their highly stable structure and their capacity to target several HKs, which confers them dual anti-virulence and anti-vancomycin resistance properties, are particularly advantageous. They can be envisioned to be used as an adjuvant in combination with vancomycin to break high-level resistance to this antibiotic. The C1LPs reported here thus represent a blueprint for engineering new HK inhibitors for the development of next generation antibiotics.

### Material and methods

### Strains, plasmids, and growth conditions

Bacterial strains used in this study are listed in Table S2. *Enterococcus* strains were cultured in M17 medium^52^ supplemented with glucose 0.5 % (GM17) at 37 °C without shaking. *Staphylococcus* strains were cultured in MH medium^53^ at 37 °C with shaking at 120 rpm. *Escherichia coli* strains were cultured in LB medium^54^ at 37 °C or 30 °C with shaking at 120 rpm. Growth was assessed by monitoring the optical density at 600 nm (OD_600_) and performed in 96-well microplates (CytoOne) in a spectrophotometer (Tecan Infinite® M Nano.) with agitation for *S. aureus* strains and without agitation for *Enterococcus* strains. Media were supplemented with chloramphenicol at 10 or 20 µg/mL (Thermo Fisher Scientific, USA), kanamycin at 15 µg/mL (Merck, USA) or tetracycline at 10 µg/mL (Serva, Germany) when necessary for mutant constructions.

All plasmids and primers used in this study are listed in Tables S3 and S4, respectively. Plasmids were replicated in *E. coli* strains. Recombinant plasmids were constructed by *in vivo* recombination using DNA fragments with overlapping ends, as previously described^55^. DNA fragments were obtained by PCR using Q5® High-Fidelity DNA Polymerase (New England BioLabs, USA). Plasmid extractions were performed with the NucleoSpin Plasmid kit (Macherey-Nagel, Germany) according to the manufacturer’s instructions. Purified plasmids were then electroporated into competent *E. faecalis* V583 cells with a Gene Pulser™ (2,25 kV, 200 Ω, 25 µF) (Biorad, USA). Mutations were introduced into the chromosome by double crossing-over of the plasmid as previously described^56^. Mutants were controlled by amplification of the corresponding loci with GoTaq DNA Polymerase (Promega, USA) and sequencing.

### Production and purification of C1LPs

The heterologous expression strain *Streptomyces coelicolor* M1146 harboring the P4H7 cosmid^23^ and the native producing strain *Streptomyces* sp. SKH 2344^57^ were used for the production of sviceucin and siamycin I, respectively. Both strains followed the same fermentation procedure. The *Streptomyces* strain was grown from a mycelium stock in 150 mL GYM medium^58^ for 4 days at 30 °C with agitation. This pre-culture was used to inoculate a 3 L fresh GYM culture, which continued growth at 30 °C for 7 days. The cell pellets were collected and extracted by 400 mL MeOH at room temperature for 4 h. After solvent evaporation, the dried crude extract was resuspended in 3 mL 80 % acetonitrile and purified by semi-preparative HPLC on a Phenomenex C18 column (Luna C18 (2), 5 µm, 100 Å, 250 x 10 mm) at 3 mL/min with UV detection at 280 nm. Eluents used were A (H2O with 0.1 % formic acid) and B (acetonitrile). For sviceucin purification, the gradient was a linear increase of 45 % to 60 % B in 10 min. For siamycin I purification, the gradient was a linear increase of 20 % to 70 % B in 30 min. The corresponding peak containing the peptide was collected. For peptide quantification, a predicted extinction coefficient at 280 nm of 11860 M^-1^.cm^-1^ for sviceucin and 7450 M^-1^.cm^-1^ for siamycin I were used.

### Determination of minimal inhibitory concentration

The MIC of vancomycin was determined by the microdilution method described in the guidelines of the Clinical and Laboratory Standards Institute (CLSI 2012). For enterococci, GM17 medium was used instead of MH medium since enterococci cannot degrade starch, the main carbohydrate of MH medium. The media were inoculated with 5.10^5^ CFU/mL. Subsequent 2-fold serial dilutions of the tested antibiotic were performed to obtain concentrations ranging from 0.5 to 512 μg/mL. Lasso peptides were then added to the wells at the desired concentration. The MIC is the lowest concentration that prevents visible *in vitro* growth.

### Extraction of muropeptides

Extraction of muropeptides was performed as previously described^59^ with modifications. Bacteria were cultured overnight in GM17 medium supplemented with 2 µg/mL of vancomycin (Merck, USA), 10 µM of sviceucin, or 2 µM of siamycin I. Bacteria were pelleted by centrifugation for 5 min at 10000 rpm and then resuspended in 1 mL of 0.1 M Tris-HCl (pH 6.8), 0.25 % SDS. Samples were boiled for 20 min in a heating block (Biolock Scientific, USA) and then centrifuged for 5 min at 10000 rpm. The pellets were washed twice with 1 mL of cold ultrapure water before sonication for 30 min (Branson 1800 Cleaner, USA). 500 µL of DNase 15 µg/mL and RNase 60 µg/mL (Sigma-Aldrich, USA) were added to each sample then incubated at 37 °C for 60 min. Samples were then treated with trypsin at 5 µg/mL (Promega, USA) and incubated at 37 °C for 2 h. Enzymes were inactivated by boiling the samples for 3 min and then washed once with ultrapure water. Pellets were resuspended in 12,5 mM sodium phosphate buffer (pH 5.5), then muropeptides concentration was determined and standardized to an OD_578_ of 3. Samples were then treated with mutanolysin (Sigma-Aldrich, USA) at 5000 U/mL and incubated at 37 °C, 150 rpm shaking, for 16 h. Samples were boiled for 3 min to inactivate the enzyme and then centrifuged. The supernatants containing the muropeptides were collected and stored for LCMS analysis.

### Quantification of muropeptides by mass spectrometry

LCMS analysis was performed on an ultra-high performance LC system (Ultimate 3000 RSLC, Thermo Scientific, USA) coupled to a high-resolution electrospray ionization-quadrupole-time of flight mass spectrometer (MaXis II ETD, Bruker Daltonics, USA). The muropeptide solution was diluted five times in MeOH before analysis. LC separation was performed on an Acclaim RSLC Polar Advantage II column (2.2 µm, 2.1 x 100 mm, Thermo Scientific, USA) with a flow-rate of 0.3 mL/min using the following gradient (eluent A: water with 0.1 % formic acid (v/v); eluent B: acetonitrile): 2 % B for 3 min, followed by a linear increase to 15 % B over 7 min and a final increase to 100 % B in 1 min. The mass range of 250-1500 in positive mode was acquired. Mass data were analyzed by Data Analysis 4.4 (Bruker Daltonics, USA). For quantification, the *m/z* corresponding to singly- and doubly-charged species of monomer, as well as doubly- and triply-charged species of dimers of muropeptides were manually searched. These were the most abundant masses observed under our experimental conditions. Subsequently, the areas of the peaks corresponding to the same muropeptide in the extracted ion chromatograms were combined for calculation.

### Analysis of gene expression by qPCR

For *gelE* gene expression measurements, bacteria were cultured in GM17 medium supplemented with the 1:1 water:methanol formulation or either sviceucin at 10 µM, or siamycin I at 1 µM. Cultures were incubated at 37 °C until reaching OD_600_ of 0.5. For *vanY* gene expression measurements, bacteria were cultured in GM17 medium supplemented with either vancomycin at 1 µg/mL or sviceucin at 10 µM and vancomycin at 1 µg/mL at 37 °C to induce the expression of *van* genes until reaching OD_600_ of 0.5. Bacteria were recovered by centrifugation at 8000 rpm for 2 min. Once the supernatant was discarded, the pellets were stored at -80 °C. Pellets were resuspended in TE buffer or ML lysis buffer, depending on kit used, and transferred in Lysis Matrix B Tubes (MP Biomedicals™, USA) before cell lysis with a FastPrep-24™ 5G (2×40 s at 6 m/s, 5 minutes on ice in-between cycles) (MP Biomedicals™, USA). After bead-beating lysis, RNA extraction was performed using Direct-Zol™ RNA Miniprep kit (Zymo Research, USA) for *gelE* gene expression measurements or NucleoSpin^Ⓡ^miRNA Miniprep kit (Macherey-Nagel, Germany) for *vanY* gene expression measurements, according to the manufacturers’ instructions. Retrotranscription was performed using QuantiTect® Reverse transcription (QIAGEN, Germany). qPCR on cDNA was performed with GoTaq® qPCR (Promega, USA). The housekeeping gene *gyrA* was used as a reference for normalization. Data were analyzed with CFX Manager software (Biorad, USA).

### Directed evolution and selection of resistant mutants

Resistant clones to sviceucin were selected by plating bacteria on GM17 agar medium supplemented with vancomycin at 32 µg/mL and sviceucin at 10 µM. Growth and MIC of vancomycin and sviceucin of the resistant clones were tested. The selection for siamycin I resistance was done by performing serial passages of bacteria in GM17 medium supplemented with vancomycin 10 µg/mL and increasing concentrations of siamycin I from 0.5 to 5 µM. Single nucleotide polymorphism (SNPs) analysis was performed on the sequenced gDNA of each selected resistant mutant.

The genomic DNA of the selected resistant mutants and the WT strain *E. faecalis* V583 were extracted using GenElute™ (Sigma-Aldrich, USA). Whole genome sequencing was then performed at the iGE3 genomics platform of the University of Geneva. The Illumina DNA prep kit was used for library preparation according to manufacturer’s specifications, with 200 ng of genomic DNA as input. Library molarity and quality was assessed with the Qubit and Tapestation using a DNA High sensitivity chip (Agilent Technologies). Libraries were sequenced on a NovaSeq 6000 Illumina sequencer with a minimum of 15 M of paired-end 100 reads per sample. The sequencing quality control was done with FastQC. The reads were mapped with the BWA v.0.7.10^60^ software to the NCBI *Enterococcus faecalis* V583 NC_004669.1 reference. After sorting and marking the PCR replicates with samtools ^61^, the alignments were screened for variants with freebayes v.0.9.21, by specifying a haploid genome.

### Nanoemulsions formulation process

The formulation of the nanoemulsions was based on the spontaneous nanoemulsification method previously described by Seguy *et al*.^33^ and transposed to lasso peptides. The present nanoemulsions were developed from anhydrous mixtures composed of excipients already described in the European Pharmacopeia (World Health Organization, 2018) and already proven to be hemocompatible on human whole blood. Monodisperse nanoemulsions could be formulated at a drug payload of 400 µg/mL for siamycin I. At these concentrations, the drug added to the anhydrous mixture during the formulation process appeared to be completely dissolved. After the addition of the aqueous phase, a homogeneous colloidal suspension was observed. The nanoemulsions were characterized in terms of granulometric and physicochemical properties (Table S5). The hydrodynamic diameter, zeta potential, and PDI were not changed relative to the blank nanoemulsions, remaining lower than 0.25, indicating the presence of a monodispersed population.

### *Galleria mellonella* infection and C1LP treatments

Bacteria were collected from overnight cultures by centrifugation, washed three times with normal saline solution (NSS), then adjusted to an OD_600_ of 0.3 corresponding to ∼10^8^ CFU/mL. 10 µL of the pre-treated bacterial suspension were injected subcutaneously into in-house reared *Galleria mellonella* larvae using a syringe pump (KD scientific, USA) in order to inoculate 10^6^ CFU into each larva or the equivalent volume of NSS as a control. To assess the anti-virulence effect of the C1LPs, *E. faecalis* V583 was grown overnight in a medium supplemented with either 10 µM sviceucin or 2 µM siamycin I. 15 larvae were used for each condition. Larvae were recorded as dead in absence of reactivity to stimuli at 24 hours p.i. To assess the C1LPs efficacy as treatment of *E. faecalis* infection, the strains V583 or V583 Δ*vanB* were grown without previous C1LP exposition. 20 larvae were used for each condition. At 1.5 and 24 hours p.i, larvae were injected with 4.76 mg of vancomycin per kilogram and/or 18 mg of siamycin I per kilogram, or the equivalent volume of nanoemulsion as a control. Larvae killing was then monitored between 1.5 and 96 h p.i. Data and statistical analysis were performed using the Kaplan-Meier R package and the logrank test, respectively.

### Proteins purification

1 L of Terrific medium^62^ equally dispatched in four 1 L baffled Erlenmeyer flasks was inoculated with an overnight culture of *E. coli* BL21DE3 possessing the plasmid pET29b(+) *vanS^127-384^* to a final of OD_600_ 0.1 (Table S2). The culture was incubated at 37 °C with agitation at 180 rpm. Induction of protein expression was achieved by adding IPTG to a final concentration of 0.1 mM when the bacterial density reached an OD_600_ of 0.5. The culture was then incubated for 20 h at 25 °C with agitation at 180 rpm. The cells were then washed twice in purification buffer (20 mM sodium phosphate buffer, 500 mM NaCl, 5 mM MgCl_2_, 10 mM imidazole (pH 7.4)). Cells were treated with lysozyme at 1 mg/mL final concentration for 30 minutes at 4 °C and lysed by sonication with the Fisherbrand Model 120 Sonic Dismembrator (Fisher scientific, USA). The total fraction recovered was centrifuged at 10000 rpm at 4 °C for 30 to 45 min and only the supernatant, representing the soluble fraction, was recovered for purification.

Protein purification was carried out on a HisTrap^TM^ FF Crude 5 mL column (Cytiva, USA) using the AKTA start device (Cytiva, USA), following the manufacturer’s recommendations. The column was equilibrated with the purification buffer and then loaded with the soluble fraction. The protein of interest was washed and eluted with an imidazole gradient of 10 mM to 500 mM in purification buffer. The purified protein was desalted and concentrated on a Vivaspin® 6 30 kDa MWCO column (Cytiva, USA) according to the manufacturer’s recommendations. Protein concentration was determined with the Pierce™ BCA Protein Assay Kit (Thermo Scientific, USA) according to the manufacturer’s recommendations and then stored at -80 °C.

### Differential scanning fluorimetry

All the experiments were performed in 96 Fast PCR Plates full skirt (Sarstedt, Germany) and in CFX96 Real-Time PCR system (BioRad, USA). Screening was conducted (Table S6) to determine the best buffer and ratios of protein and SYPRO® Orange Protein Gel Stain (Sigma-Aldrich, USA). For the experiments, each well contained 25 µL of 10 µM VanS solution, 1/450 X of SYPRO® Orange Protein Gel Stain and lasso peptides at various concentrations: siamycin I – 6 to 90 µM; sviceucin – 12 to 180 µM; microcin J25 – 25 to 200 µM. VanS, SYPRO® Orange and lasso peptides were diluted in 20 mM potassium phosphate buffer (pH 7.5, Table S6). Each condition was replicated three times per plate and each plate was replicated three times. CFX Manager was used to recover data from runs. The Tm of each well was determined on the Melt Curve panel with a negative peak type. Tm means were calculated for each condition before model analysis. ΔT was calculated from the subtraction of the Tm of the protein with the buffer alone and the Tm with the lasso peptides. All analyses were carried out as previously described^63^.

### Autophosphorylation assays for VanS

The phosphorylation assay followed reported procedure with some modifications^38^. A standard reaction of 200 µL contained 50 mM Tris-HCl (pH 7.8), 100 mM NaCl, 2 µM purified C-His_6_-VanS^127-384^, 0-100 µM of respective lasso peptide, 5 mM ATP, 10 mM MgCl_2_, 0.5 mM NADH, 2 mM phospho(enol)pyruvic acid (PEP), 18 U of L-lactate dehydrogenase (LDH, Roche, Switzeland) and 9 U of pyruvate kinase from rabbit muscle (PK, Sigma-Aldrich, USA). The peptide was pre-incubated with the kinase at room temperature for 10 min followed by the addition of PEP/LDH/PK and incubation at 37 °C for 3 min. The reaction was allowed to start by adding ATP and NADH, and monitored continuously at 340 nm at 37 °C by a microplate reader (POLARstar OMEGA, BMG Labtech, Germany). Each measurement was performed in triplicate, and at least three independent experiments were repeated. Nonlinear regression data fitting and IC_50_ calculation were performed with GraphPad Prism 10.

## Acknowledgements

The funding supports from the ANR (Agence Nationale de la Recherche) (PRC project no. ANR-19-CE18-0026-01) and the SATT-Lutech are acknowledged. A.B and B.B thank the Normandy region and the French research ministry for the Ph.D funding. DNA sequencing was performed at the iGE3 genomics platform of the University of Geneva (http://www.ige3.unige.ch/genomicsplatform.php) and we thank Mylene Docquier for her expertise. We also thank Nicholas J. Harmer for his help in the DSF data analysis process.

## Supplementary data

**Figure S1.**
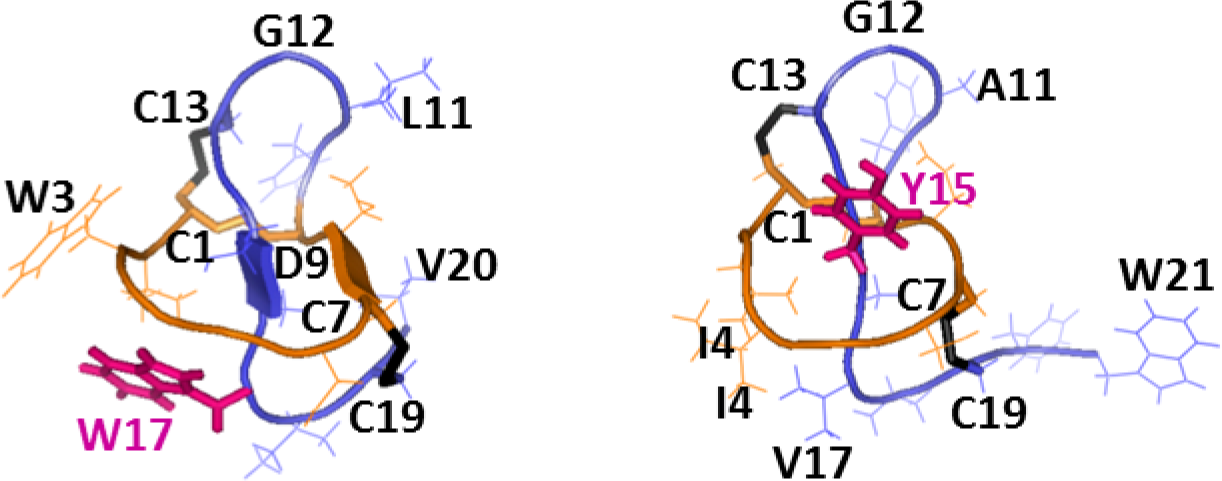
Three-dimensional structures of class I lasso peptides. Left: sviceucin^2^ (PDB: 2LS1); Right: RP-71955^3^ (PDB: 1RPC) which is a close homolog of siamycin I (primary sequence: CLGIGSCNDFAGCGYAVVCFW).

**Figure S2.**
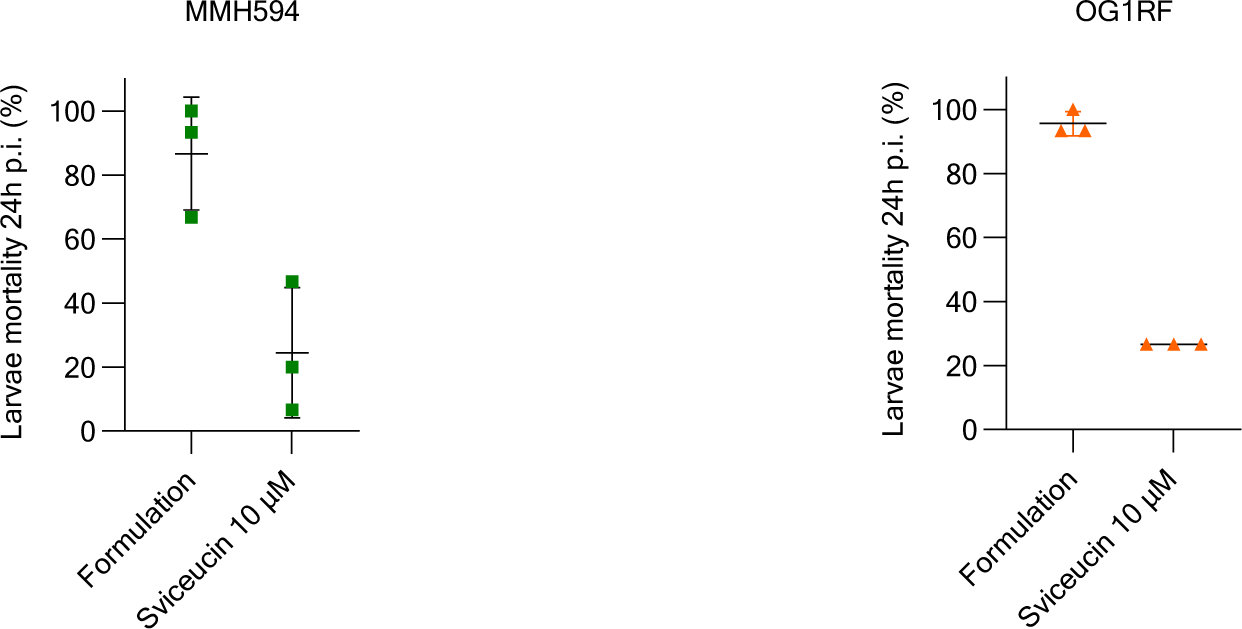
Survival of *Galleria mellonella* larvae infected with *E. faecalis* MMH594 or OG1RF treated with sviceucin.

**Figure S3.**
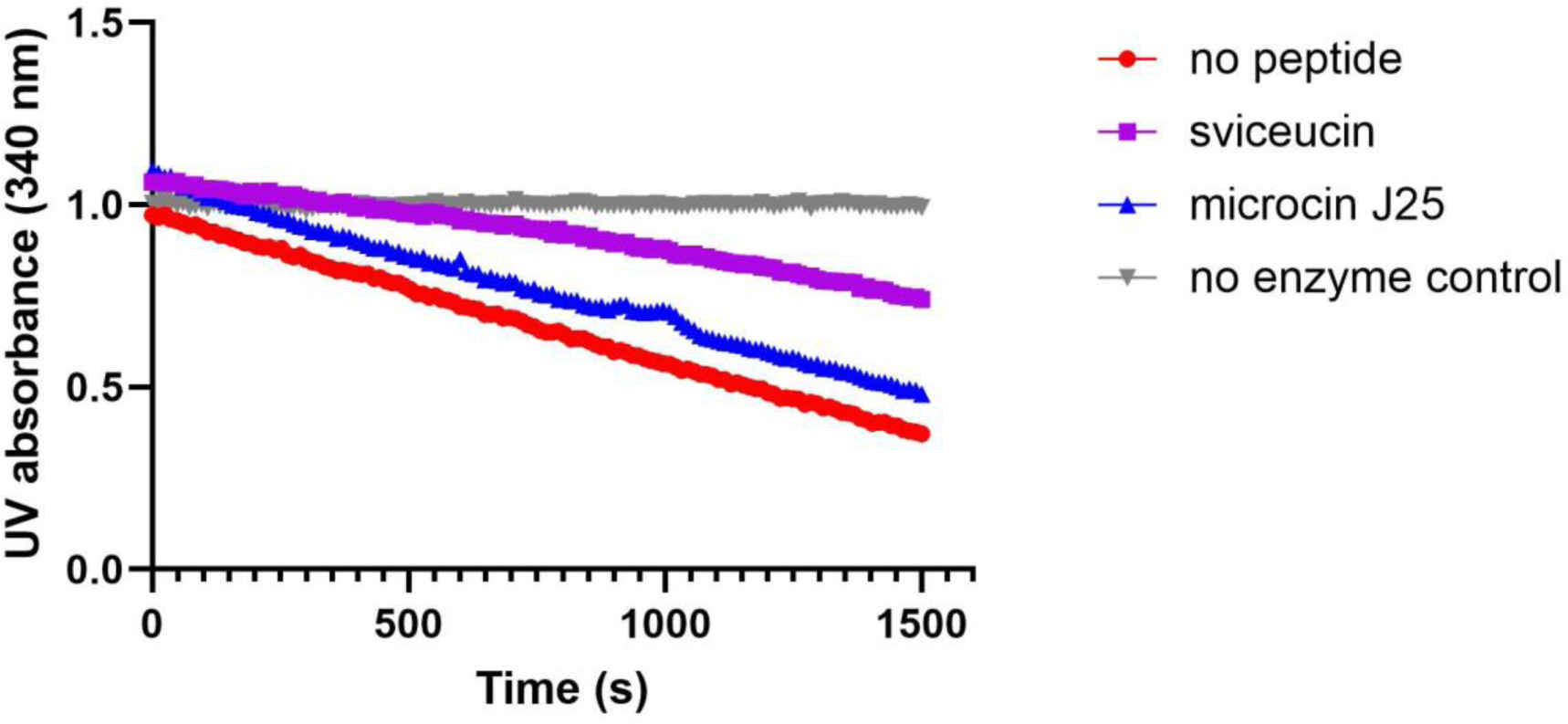
Autophosphorylation activity of VanS^122-384^ in the presence of lasso peptides. A continuous assay with coupling enzymes was performed as previously described^38^. Reaction rate was reflected by the consumption of NADH monitored by UV spectroscopy at 340 nm.

**Figure S4.**
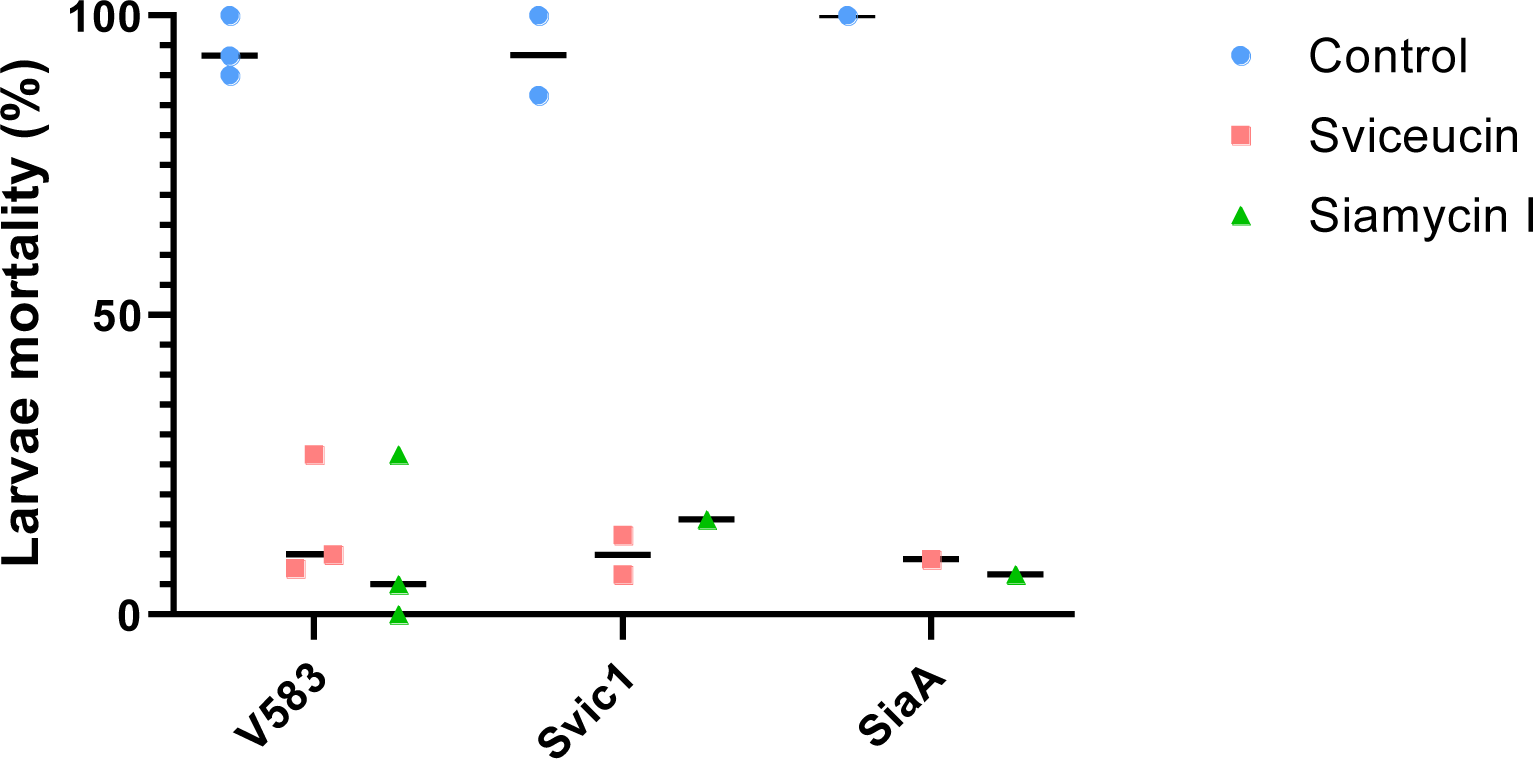
Anti-virulence activity of sviceucin and siamycin I on V583, Svic1 and SiaA. Mortality of *Galleria mellonella* larvae 24 h post-infection (p.i.) with *E. faecalis* V583, mutant Svic1 and mutant SiaA, pre-treated with either siamycin I (2 µM), sviceucin (10 µM) or formulation (control).

**Table S1.** Characteristics of the spontaneous mutants resistant to lasso peptides and vancomycin combinations.

| Clone no. | Isolation method | Position in v583 genome | V583 reference | Mutation | Mutations consequences | Vancomycin MIC (µg/ml) | Vancomycin + svceucin MIC (µg/ml) | Vancomycin + siamycin i MIC (µg/ml) |
| --- | --- | --- | --- | --- | --- | --- | --- | --- |
| V583 | n.a. | n.a. | n.a. | n.a. | parental strain | 32 | 2 | 2 |
| Svic1 | 5*10<sup>8</sup> plating on vancomycin (32 µg/ml) and svceucin (10 µM) | 2216922 | C | A | VanSB S402I | 128 | 128 | 2 |
| Svic2 |  | 2527311 | C | A | non sense mutation in *atpA* | 256 | 256 | 2 |
| Svic3 |  | 2529182 | C | A | non sense mutation in *atpF* | 256 | 256 | 2 |
| Svic4 |  | 2525799 | C | T | *atpD* RBS modification | 256 | 256 | 2 |
| SiaA | serial passages | 2216933 | A | C | VanSB D398E | 128 | 128 | 32 |
| SiaB |  | 1690453 | T | A | CcpA D121V | 64 | 32 | 32 |
| SiaC |  | 1690372 | G | A | CcpA A148V | 64 | 32 | 32 |
| SiaD |  | 1690550 | C | G | CcpA A89P | 64 | 32 | 32 |
| SiaE |  | 1690580 | A | deletion | frameshift in *ccpA* | 32 | 16 | 16 |

**Table S2.**
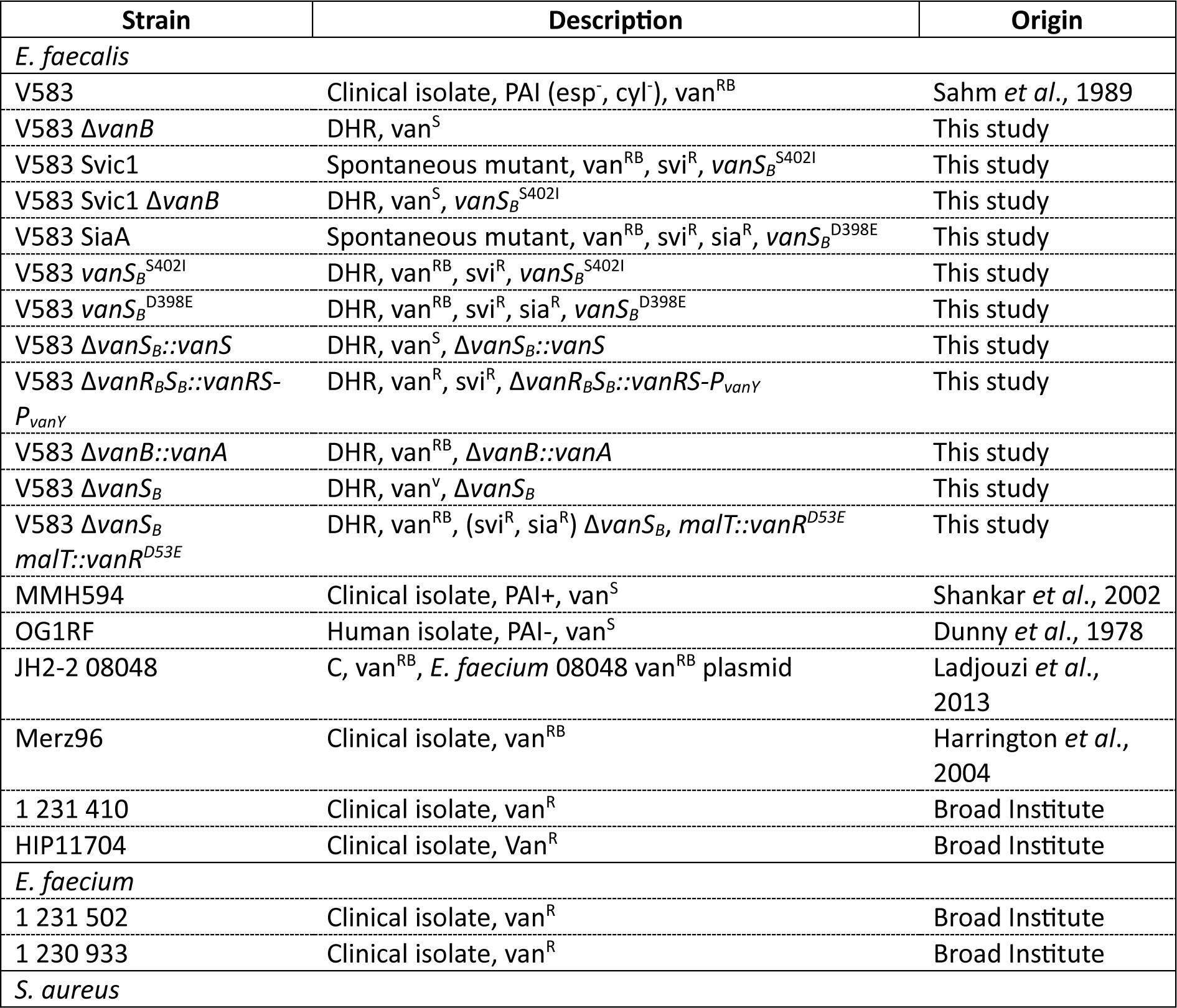
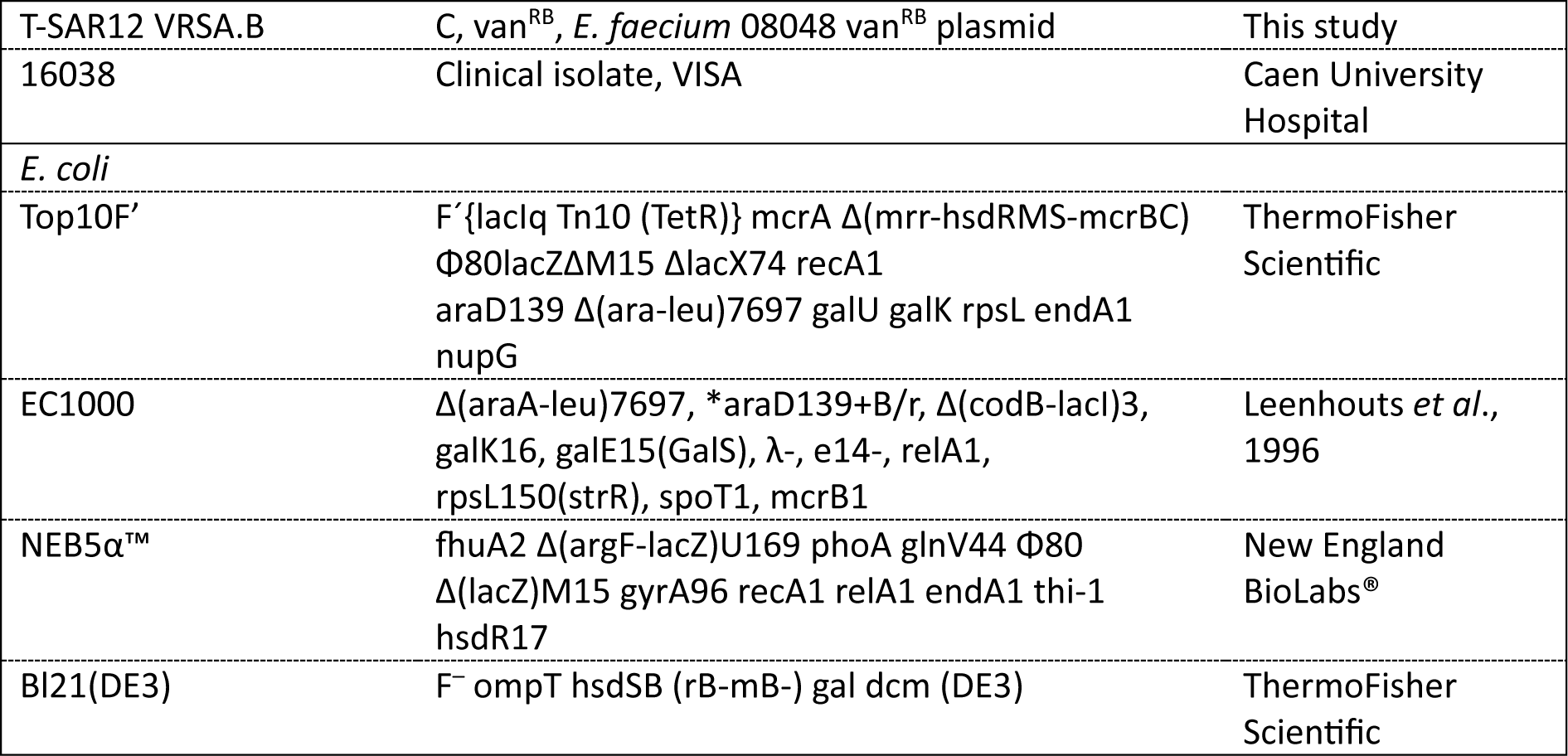
Strains used for this study. DHR = Double Homologous Recombination, C = Conjugation, R = Resistant, S = Sensitive, V = Variable

**Table S3.** Plasmids used for this study.

| Plasmid | Description | Origin |
| --- | --- | --- |
| pEBM2 | oriT, *repA* TS, lac promoter, lac operator,<br>*lacZ*, *thyA* *, Cm<sup>R</sup><br>For *E. coli*: replicative<br>For *E. faecalis*: replicative at 30°C and<br>integrative at 37°C | Lab collection |
| pEBM2 Δ*vanS<sub>B</sub>* | pEBM2-(*vanR<sub>B</sub>*-*VanY<sub>B</sub>*) | This study |
| pEBM2 Δ*vanS<sub>B</sub>::vanS* | pEBM2-(*vanR<sub>B</sub>*-*VanS*-*VanY<sub>B</sub>*) | This study |
| pEBM2 Δ*vanR<sub>B</sub>S<sub>B</sub>::vanRS-P<sub>vanH</sub>* | pEBM2-(*vanR<sub>B</sub>* upstream-*vanR*-*vanS*-<br>*P<sub>vanH</sub>*-*vanY<sub>B</sub>*) | This study |
| pEBM2 *maltT::P<sub>hup</sub>-vanR<sup>D53E</sup>* | pEBM2-(start *malt*-*P<sub>hup</sub>-vanR<sup>D53E</sup>*-end<br>*malt*) | This study |
| pAS222 | Tet<sup>R</sup>, Amp<sup>R</sup>, *repA*-pG+host4<br>For *E. coli* : replicative<br>For *E. faecalis* : replicative at 30°C and<br>integrative at 37°C | Jönsson *et al.*, 2009 |
| PAS222 *vanS<sub>B</sub><sup>S402I</sup>* | pAS222-(*vanR<sub>B</sub>*-*vanS<sub>B</sub><sup>S402I</sup>*-*vanY<sub>B</sub>*) | This study |
| PAS222 *vanS<sub>B</sub><sup>D398E</sup>* | pAS222-(*vanR<sub>B</sub>*-*vanS<sub>B</sub><sup>D398E</sup>*-*vanY<sub>B</sub>*) | This study |
| pWS3 | pG<sup>+</sup>host9 derivative, spectinomycin<sup>R</sup>,<br>erythromycin<sup>S</sup><br>Suicide vector in *E. faecalis* | Zhang *et al.*, 2011 |
| pWS3 Δ*vanB* | pWS3-(*vanHB*-*vanXB*) | This study |
| pWS3 Δ*vanB::vanA* | pWS3-(*vanHB*-*vanA*-*vanXB*) | This study |
| pET29b(+) | Kan<sup>R</sup>, T7lac, His-tag, S-tag, cleavage of<br>tags by thrombin | (Merck ?) |
| pET29b(+) *vanS<sub>127-384</sub>* | Heterologous production in *E. coli* | This study |
| pET29b(+) *croS<sub>133-393</sub>* | Heterologous production in *E. coli* | This study |

**Table S4.**
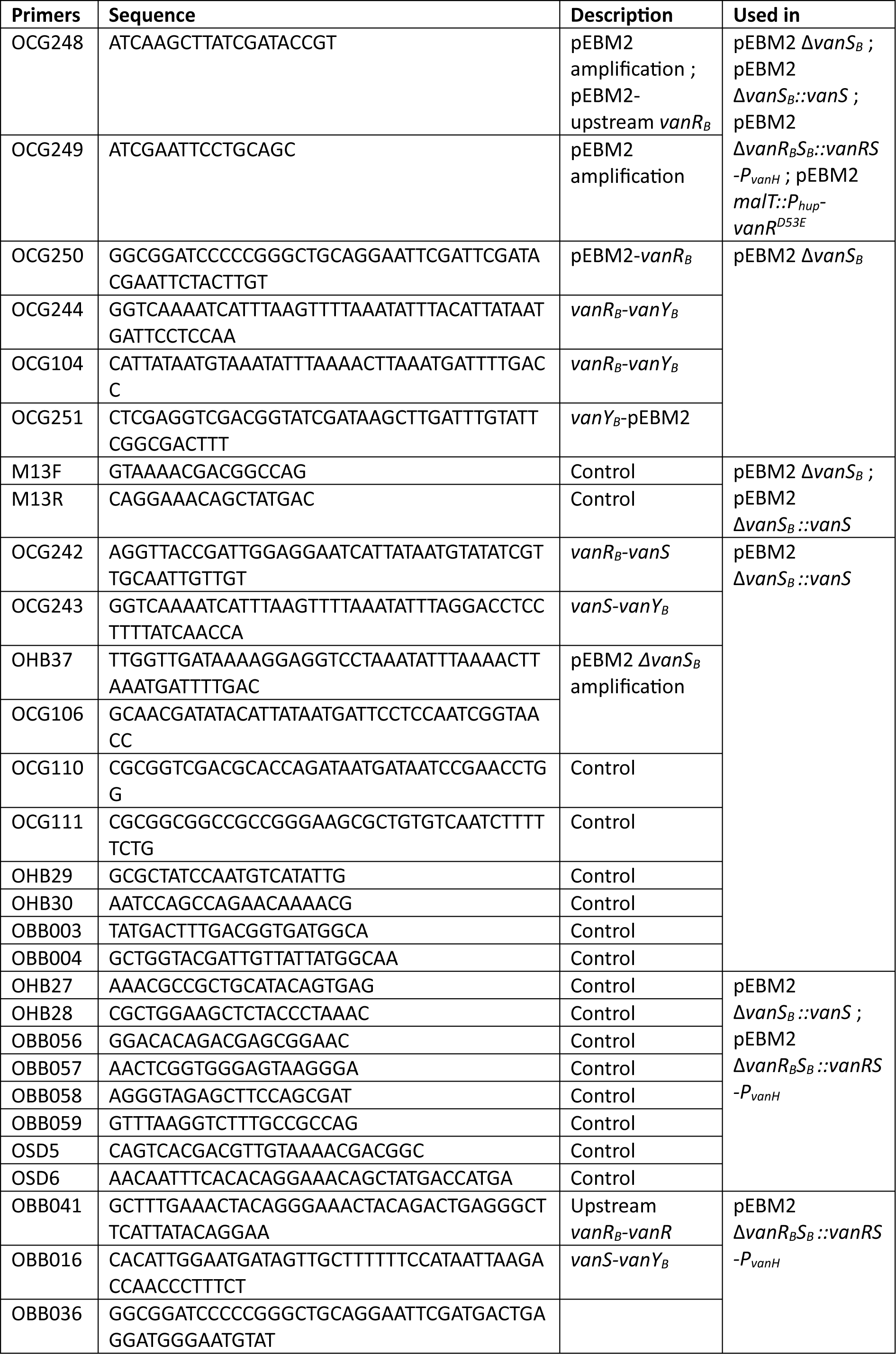
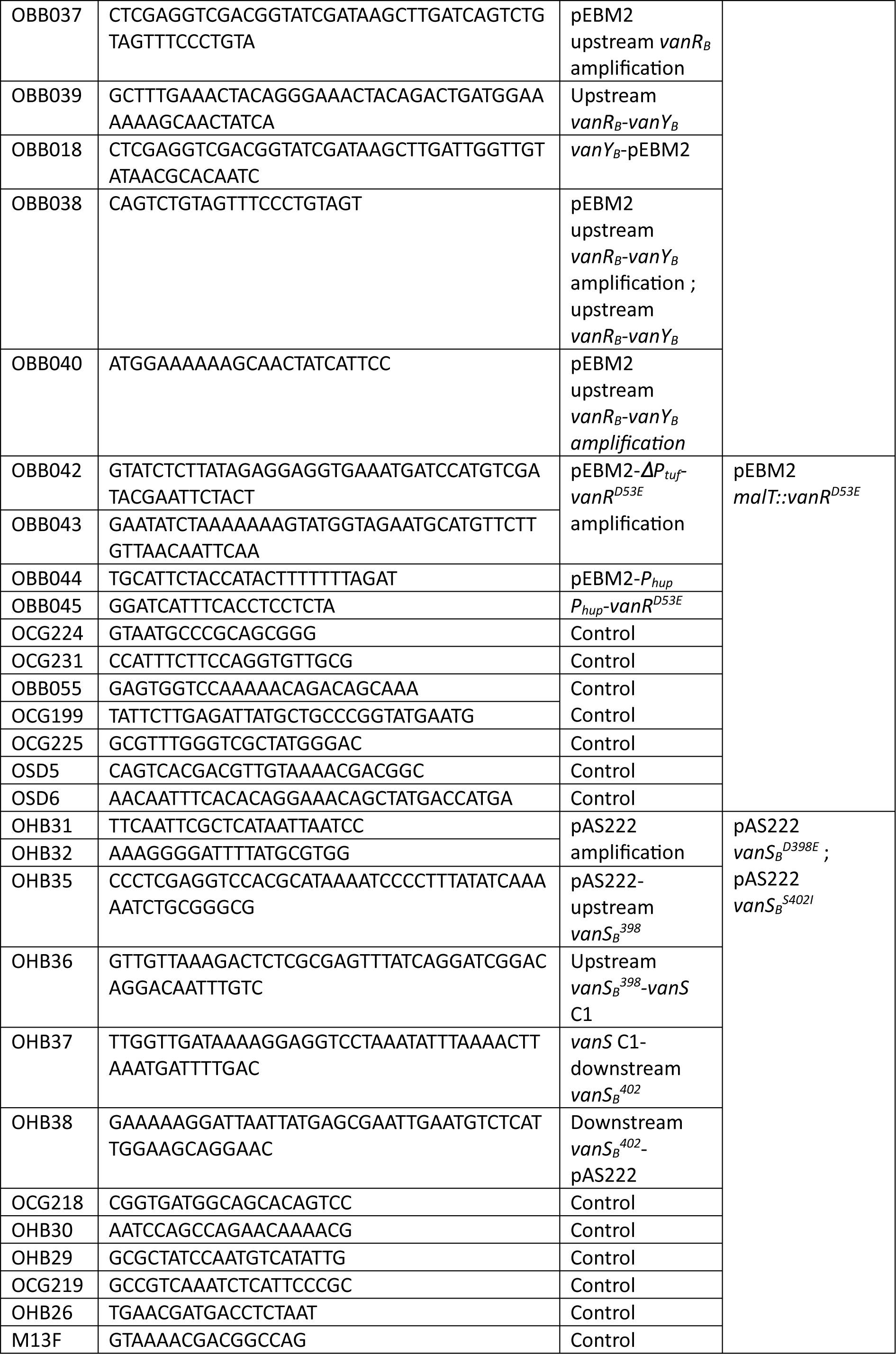
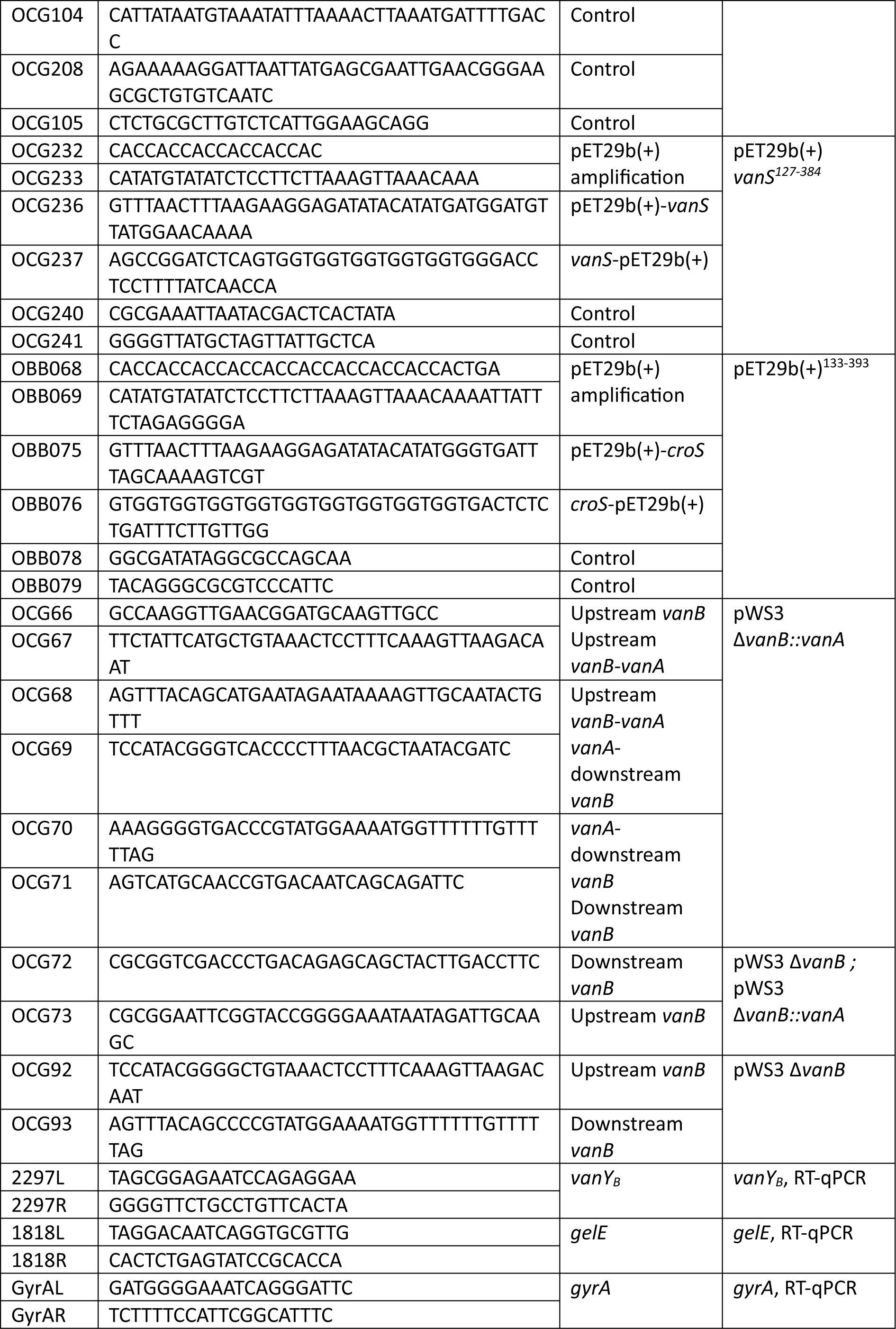
Primers used for this study.

**Table S5.** Physicochemical characteristics of the uncharged and C1LP containing nanoemulsions.

|  | **Z-average diameter (nm)** | **PDI** | **Zeta potential (mV)** | **Vancomycin MIC on *E. faecalis* V583** |
| --- | --- | --- | --- | --- |
| Blank NEs | 54 ± 2 | 0.21 ± 0.02 | -16 ± 4 | 32 |
| Sviceucine-NEs | 52 ± 1 | 0.17 ± 0.03 | -15 ± 5 | 4 |
| Siamycin-NEs | 51 ± 1 | 0.17 ± 0.01 | -17 ± 1 | 4 |

**Table S6.** Buffers screened for DSF.

| **Buffers** | **Final concentration** |
| --- | --- |
| Tris pH 7.5 | Tris 50 mM |
|  | Glycerol 15 % |
| Tris NaCl pH 7.5 | Tris 50 mM |
|  | Glycerol 15 % |
|  | NaCl 150 mM |
| Tris MgCl<sub>2</sub> pH 7.5 | Tris 50 mM |
|  | Glycerol 15 % |
|  | MgCl<sub>2</sub> 5 mM |
| Tris NaCl/MgCl<sub>2</sub> pH 7.5 | Tris 50 mM |
|  | Glycerol 15 % |
|  | NaCl 150 mM |
|  | MgCl<sub>2</sub> 5 mM |
| Potassium phosphate pH 7.5 | Potassium phosphate 20 mM |
|  | Glycerol 10 % |
| Potassium phosphate NaCl pH 7.5 | Potassium phosphate 20 mM |
|  | Glycerol 10 % |
|  | NaCl 500 mM |
| Potassium phosphate MgCl<sub>2</sub> pH 7.5 | Potassium phosphate 20 mM |
|  | Glycerol 10 % |
|  | MgCl<sub>2</sub> 5 mM |
| Potassium phosphate NaCl/MgCl<sub>2</sub> pH 7.5 | Potassium phosphate 20 mM |
|  | Glycerol 10 % |
|  | NaCl 500 mM |
|  | MgCl<sub>2</sub> 5 mM |
| Sodium phosphate pH 7.5 | Sodium phosphate 20 mM |
|  | Glycerol 10 % |
| Sodium phosphate NaCl pH 7.5 | Sodium phosphate 20 mM |
|  | Glycerol 10 % |
|  | NaCl 100 mM |
| Sodium Citrate pH 7.5 | Sodium citrate 50 mM |
|  | Glycerol 10 % |
| HEPES pH 7.5 | HEPES 50 mM |
|  | Glycérol 10 % |
| HEPES NaCl pH 7.5 | HEPES 50 mM |
|  | Glycérol 10 % |
|  | NaCl 100 mM |
| HEPES MgCl<sub>2</sub> pH 7.5 | HEPES 50 mM |
|  | Glycérol 10 % |
|  | MgCl<sub>2</sub> 5 mM |
| HEPES NaCl/MgCl<sub>2</sub> pH 7.5 | HEPES 50 mM |
|  | Glycérol 10 % |
|  | NaCl 100 mM |
|  | MgCl<sub>2</sub> 5 mM |
| PIPES pH 7.5 | PIPES 50 mM |
|  | Glycerol 10 % |
| MOPS pH 7.5 | MOPS 50 mM |
|  | Glycerol 10 % |
| MOPS NaCl pH 7.5 | MOPS 50 mM |
|  | Glycerol 10 % |
|  | NaCl 10 mM |
| MOPS MgCl<sub>2</sub> pH 7.5 | MOPS 50 mM |
|  | Glycerol 10% |
|  | MgCl<sub>2</sub> 5 mM |
| MOPS NaCl/MgCl<sub>2</sub> pH 7.5 | MOPS 50 mM |
|  | Glycerol 10 % |
|  | NaCl 10 mM |
|  | MgCl<sub>2</sub> 5 mM |

## Notes

### Competing Interest Statement

The authors have declared no competing interest.

